# Hormone Receptor-status Prediction in Breast Cancer Using Gene Expression Profiles and Their Macroscopic Landscape

**DOI:** 10.1101/2020.03.29.014050

**Authors:** Seokhyun Yoon, Hye Sung Won, Keunsoo Kang, Kexin Qiu, Woong June Park, Yoon Ho Ko

## Abstract

The cost of next-generation sequencing technologies is rapidly declining, making RNA-seq-based gene expression profiling (GEP) an affordable technique for predicting receptor expression status and intrinsic subtypes in breast cancer (BRCA) patients. Based on the expression levels of co-expressed genes, GEP-based receptor-status prediction can classify clinical subtypes more accurately than can immunohistochemistry (IHC). Using data from the cancer genome atlas TCGA BRCA and METABRIC datasets, we identified common predictor genes found in both datasets and performed receptor-status prediction based on these genes. By assessing the survival outcomes of patients classified using GEP- or IHC-based receptor status, we compared the prognostic value of the two methods. We found that GEP-based HR prediction provided higher concordance with the intrinsic subtypes and a stronger association with treatment outcomes than did IHC-based hormone receptor (HR) status. GEP-based prediction improved the identification of patients who could benefit from hormone therapy, even in patients with non-luminal BRCA. We also confirmed that non-matching subgroup classification affected the survival of BRCA patients and that this could be largely overcome by GEP-based receptor-status prediction. In conclusion, GEP-based prediction provides more reliable classification of HR status, improving therapeutic decision making for breast cancer patients.

## 1. Introduction

Breast cancer (BRCA) is a highly heterogeneous disease that involves several complex molecular networks [1–7]. BRCA can be classified into different subtypes that have distinct clinical behaviors and prognoses and that require different treatment strategies, including targeted therapy and hormone therapy. Therefore, accurate classification of BRCA subtypes is crucial for personalized disease management and for improving patient outcomes [8,9]. Currently, therapeutic decision making in BRCA is based on the expression status of three receptors: estrogen receptor (ER), progesterone receptor (PR), and human epidermal growth factor receptor 2 (HER2) [10, 11]. Although ER, PR, and HER2 status is traditionally determined by immunohistochemistry (IHC), with the advent of high-throughput technologies for gene expression analysis, new molecular subtypes of BRCA have been described. These include luminal A, luminal B, HER2-enriched, basal-like, and normal-like breast tumors [12–14]. The clinical significance of these intrinsic BRCA subtypes has been highlighted by their ability to predict treatment response and prognosis [4–7, 15–21]; hence their use in clinical practice has increased over recent years. Currently, several gene-signature tests based on microarray or quantitative real-time PCR (qRT-PCR) are commercially available [9, 22, 23].

The clinicopathological surrogate definitions of the intrinsic BRCA subtypes were endorsed by the 2013 St. Gallen Consensus Recommendations [24]. Luminal A BRCA is hormone receptor (HR) positive, HER2 negative, and expresses low levels of the protein Ki-67. Luminal B BRCA is HR positive and either HER2 positive or HER2 negative, with high levels of Ki-67. The HER2-enriched subtype is HR negative and HER2 positive, and the basal-like subtype is HR negative and HER2 negative (triple-negative BRCA) [25 – 27]. Although the expression profiles and clinical features of the four intrinsic BRCA subtypes have been extensively studied in the last few years, discordance has been reported between IHC-based clinical subtypes and intrinsic subtypes in approximately 20–50% of cases [18, 28, 29]. This discordance might be due to intratumoral heterogeneity, the coexistence of cells with different subtypes in the same tumor, as well as measurement inaccuracies in subtype profilers, IHC analysis for ER/PR status, and fluorescence in situ hybridization (FISH) analysis for HER2 status. These inconsistencies could result in administration of the wrong treatment, subsequently leading to poor survival [30]. Therefore, accurate identification of receptor status or the intrinsic BRCA subtype is of high clinical importance.

Recently, multi-omics technologies [31], miRNA profiling [32] and principle component analysis-based iterative PAM50 subtyping [33] have helped to improve the accuracy of BRCA subtype classification. However, inconsistencies due to measurement noise remain a challenge in this classification, especially for tumors with receptor expression levels at the boundary between positive and negative [33]. With the development of next-generation sequencing (NGS) technologies, the cost of gene expression profiling (GEP) based on RNA-seq is rapidly decreasing, making it possible to characterize several clinical and molecular features concurrently using RNA-seq-based GEP at a very low cost [34, 35]. Prediction of the intrinsic subtype and receptor status (ER, PR, or HER2) in BRCA using RNA-seq-based GEP would increase the clinical usefulness of RNA-seq technologies in BRCA. In this study, we assessed whether variations in gene expression are reflected in the expression of related genes and whether these changes can be identified by GEP to provide more reliable prediction of the status of the three receptors, thereby improving therapeutic decision making.

## 2. Results

### 2.1. Identification of predictor genes

In this study, IHC-based characterization of receptor status in BRCA was refined by using co-expressed predictor genes. First, predictor genes were identified; seven genes were selected for ER status prediction, six for PR, and four for HER2 (Table 1). As expected, the *ESR1*, *PGR*, and *ERBB2* genes, which encode the ER, PR, and HER2 proteins, respectively, were among the predictor genes. Model training and receptor-status prediction were then performed using the selected genes. The mismatch rate reported in Table 1 is the percentage of cases in which the IHC-based status differed from the predicted status. Among the predictor genes, *TFF1* and *NAT1* were included in an eighteen-gene set previously reported to predict sensitivity to hormone therapy [36].

**Table 1.** Summary of mismatch rates and predictor genes for ER, PR, and HER2 status prediction.

| Item | Mismatch rate [%]* |  | Predictor genes |
| --- | --- | --- | --- |
|  | TCGA | METABRIC |  |
| ER | 6.28 | 6.26 | *ESR1*, *AGR3*, *C1orf64*, *C4orf7*, *CLEC3A*, *SOX11*, *TFF1* |
| PR | 11.43 | 5.54 | *PGR*, *AGR3*, *ESR1*, *NAT1*, *PVALB*, *S100A7* |
| HER2 | 11.85 | 5.17 | *ERBB2*, *CPB1*, *GSTT1*, *PROM1* |

\* Between the IHC-based and the predicted receptor status

### 2.2. Macroscopic landscape

Figure 1 shows uniform manifold approximation and projection (UMAP) plots [38] for receptor status in the TCGA BRCA cohort. Each point represents a sample; the color of the spots corresponds to the (a) subtype (PAM50 class), (b) ER status, (c) PR status, and (d) HER2 status of the sample. Receptor status (ER, PR, or HER2) was provided in the original clinical data based on IHC. The expression of 100 genes selected by LASSO was used to obtain the two-dimensional UMAP projection. The luminal A and B subtypes were mostly HER2− and either ER+ or PR+. However, a small percentage of the luminal A and B subtypes exhibited ER−, PR−, and HER2+. Some patients with HER2-enriched or basal-like subtype BRCA also showed some level of discordance, as some HER2-enriched and basal-like subtype samples were ER+ or PR+. Although most HER2+ and HER2-enriched subtype samples overlapped, some HER2-enriched subtype samples were found to be HER2− BRCA that exhibited basal-like subtype features. As only eight patients exhibited normal-like subtype BRCA in the TCGA dataset, they were not considered in our analyses.

**Figure 1.**
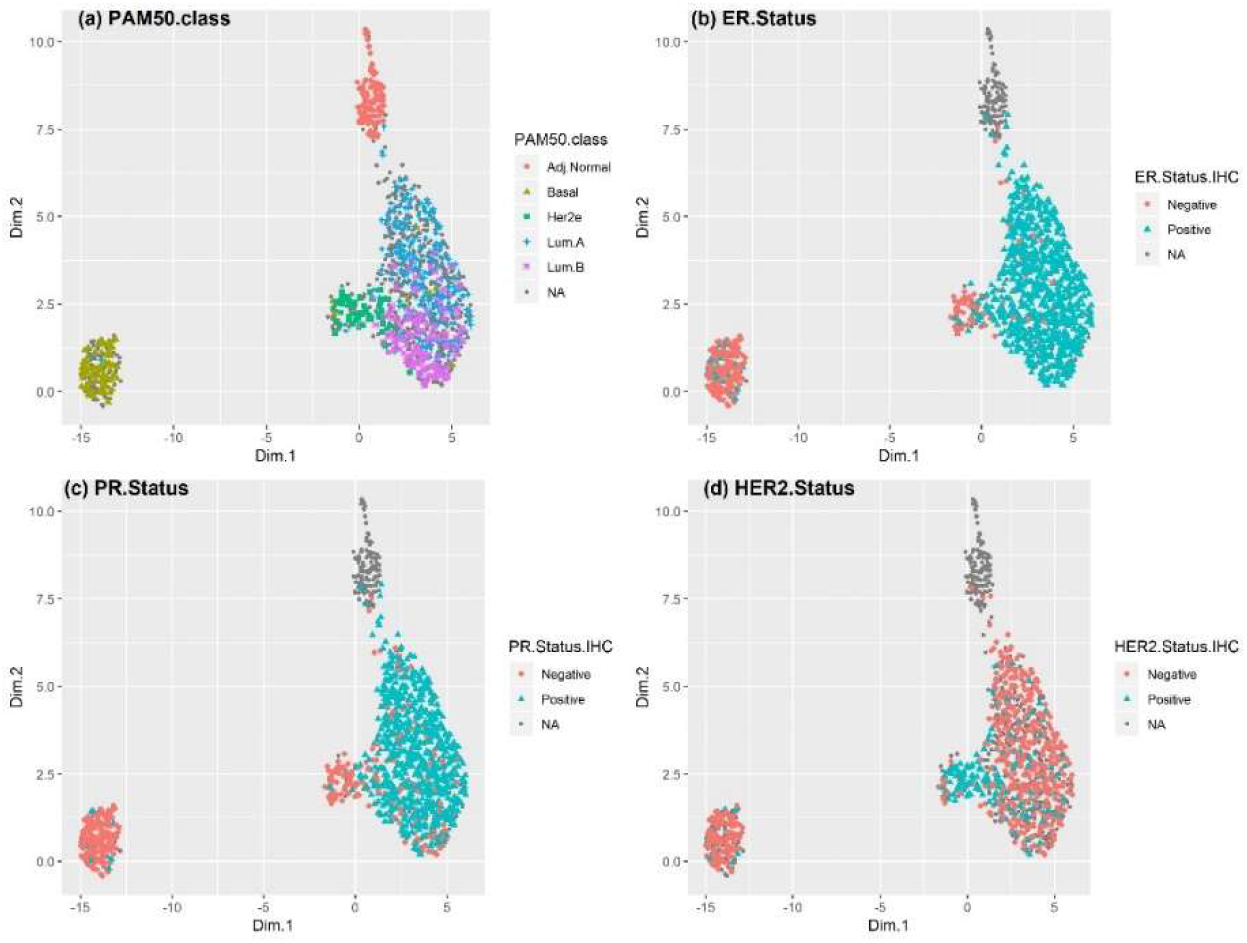
UMAP plot showing the receptor status in the TCGA BRCA cohort. The tumor subtype, as well as the status of ER, PR, and HER2, were based on the available clinical data. Gray points are samples with no available clinical information. A small percentage of the luminal A and B subtypes were ER−/PR− and HER2+. Such discordances were also observed in some BRCA patients with the HER2-enriched and basal-like subtypes. Although most HER2+ and HER2-enriched subtype samples overlapped, some HER2-enriched subtype samples were found to be HER2− BRCA and to exhibit basal-like subtype features. Some samples were ER+ and/or PR+, representing a luminal subtype.

On the other hand, the HER2-enriched subtype samples were ER+ and/or PR+, representing a luminal subtype. The UMAP plot of the METABRIC dataset revealed a similar macroscopic landscape (Supplementary Figure 1). Considering that the distance between samples (points) in the UMAP projection is only an approximation of the relative distance in their gene expression profiles and that the receptor status was not clearly defined for all samples, Figure 1 implies that IHC/FISH-based characterization of receptor status might result in inaccuracies in BRCA subtype classification.

Figure 2 shows the same UMAP plot based on the predicted values obtained by the linear classifiers. Compared with IHC-based receptor-status characterization, the predicted status was more consistent with the intrinsic BRCA subtype classification, especially for the basal-like and luminal subtypes. Most of the luminal subtypes were ER+ and PR+, and the numbers of ER+ or PR+ samples in the basal-like subtype were much smaller than after IHC-based status characterization. The UMAP plot for the METABRIC dataset based on the predicted receptor status (Supplemental Figure 2) led to the same conclusions, except for PR status, which was not IHC-based in the METABRIC dataset.

**Figure 2.**
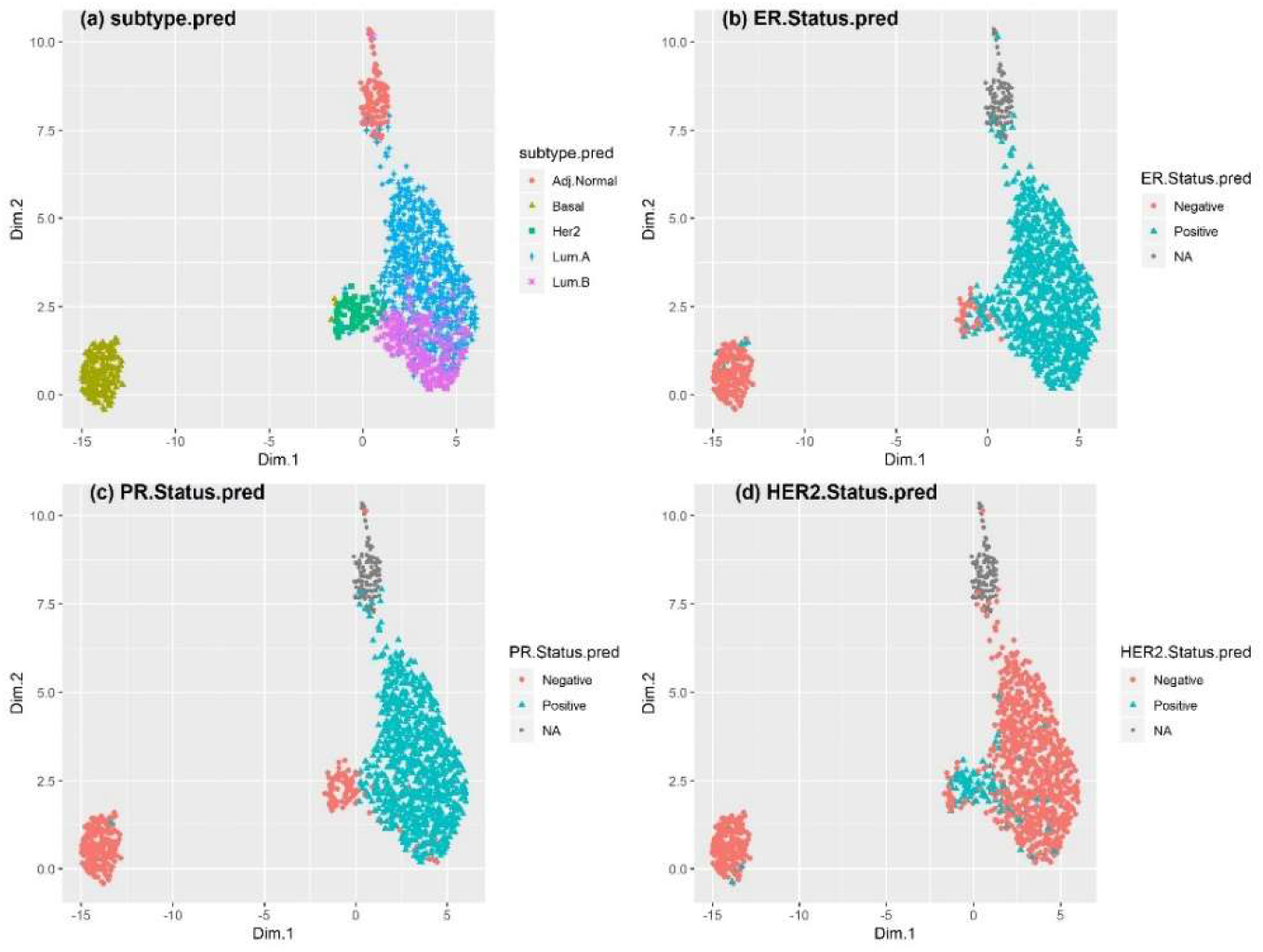
UMAP plot showing GEP-based receptor status in the TCGA BRCA cohort. GEP-based prediction was used to determine the subtype, as well as the status of ER, PR, and HER2. Compared to the case with IHC-based approaches, the predicted status of ER, PR, and HER2 was mostly in accordance with the corresponding pattern of receptor status for basal-like, luminal A, and luminal B. In contrast, HER2-enriched subtype tumors were highly heterogeneous.

### 2.3. GEP-based receptor-status prediction is reliable for the luminal and basal-like subtypes

To quantify discordance between the intrinsic subtype and the clinical subtype defined by HR and HER2 status, for each intrinsic subtype, we compared the numbers of positive and negative instances of HR and HER2 status based on IHC with the numbers obtained using GEP-based prediction in the TCGA and METABRIC datasets (Table 2). The rates of discordance for the basal-like, luminal A, and luminal B subtypes were lower using GEP-based prediction than using IHC-based status characterization. Specifically, most samples of the luminal A and B subtypes were characterized as HR+ by GEP-based prediction (except for two samples in the TCGA BRCA cohort), while some luminal A and luminal B BRCA samples were characterized as HR- based on IHC. In BRCA patients with the basal-like subtype, a smaller percentage of tumors was determined to be HR+ using GEP-based prediction (10% in TCGA and 13% in METABRIC) than when using IHC-based characterization (17% in TCGA and 20% in METABRIC).

**Table 2.** HR and HER2 status for each intrinsic subtype as determined by (a) IHC- and (b) GEP-based prediction. Patients with no available IHC-based receptor status were excluded.

| Dataset | Subtype | (a) IHC-based characterization |  | (b) GEP-based prediction |  |
| --- | --- | --- | --- | --- | --- |
|  |  | HR+/- | HER2+/- | HR+/- | HER2+/- |
| TCGA | Luminal A | 222 / 4 | 24 / 130 | 229 / 2 | 4 / 227 |
|  | Luminal B | 126 / 1 | 22 / 69 | 127 / 0 | 8 / 119 |
|  | Basal-like | 16 / 78 | 6 / 59 | 10 / 87 | 2 / 95 |
|  | HER2-enriched | 32 / 24 | 40 / 10 | 44 / 14 | 39 / 19 |
| METABRIC | Luminal A | 680 / 6 | 19 / 283 | 696 / 0 | 19 / 677 |
|  | Luminal B | 465 / 1 | 23 / 171 | 474 / 0 | 29 / 445 |
|  | Basal-like | 61 / 243 | 14 / 118 | 40 / 268 | 24 / 284 |
|  | HER2-enriched | 119 / 111 | 50 / 34 | 125 / 111 | 119 / 117 |
|  | Normal-like | 161 / 21 | 11 / 51 | 165 / 19 | 11 / 173 |

On the other hand, considerable discordance was observed in the receptor status of HER2-enriched subtype BRCA patients using both IHC-based characterization and GEP-based prediction. Only 37% and 23% of patients with HER2-enriched subtype BRCA were HR-/HER2+ in the METABRIC and TCGA datasets, respectively. Furthermore, 17% and 18% of tumors were triple negative, and 25% and 9% were luminal-like (HR+ and HER2-) in the METABRIC and TCGA datasets, respectively. Similar findings were obtained for IHC-based characterization of HR and HER2 status.

In summary, GEP-based prediction was more concordant with the typical receptor-status pattern of the intrinsic subtypes of patients with the basal-like, luminal A, and luminal B subtypes. However, this does not necessarily mean that receptor-status prediction based on GEP is more accurate than IHC-based characterization. The only way to verify the accuracy of the status predictions is to assess the differences in clinical outcomes among the different clinical subtypes defined by the status of the three receptors.

### 2.4. GEP-based receptor-status prediction is reliable for the luminal and basal-like subtypes

To verify the accuracy of the receptor-status predictions, survival outcomes for various combinations of HR and HER2 status were compared. The significance of the prognostic value of the predicted and IHC-characterized HR and HER2 status was compared. Separate survival analyses were performed in the following four patient groups:

a. HR+ (either ER+ or PR+) group: This group benefited from hormone therapy. According to the stage and clinical characteristics, these patients often received a combination of hormone therapy and chemotherapy. For survival analysis, the patients in this group were stratified based on administration of hormone therapy.
b. Hormone therapy group: To confirm the benefit of hormone therapy for HR+ patients, only those who received hormone therapy, with or without chemotherapy, were selected, and the survival of HR+ patients was compared to that of HR– patients.
c. HR+/non-luminal subtype group: As shown in Table 2, there were small percentages of HR+ patients among patients with the HER2-enriched and basal-like subtypes. Hence, we assessed whether BRCA patients with the HR+ non-luminal subtype benefited from hormone therapy.
d. HER2+ group: BRCA patients with the HER2+ subtype benefited from anti-HER2 targeted molecular therapy (TMT). We assessed the survival of HER2+ BRCA patients based on TMT. As no information regarding TMT was available in the METABRIC dataset, this analysis was performed only for the TCGA BRCA cohort.

Among patients in the TCGA BRCA cohort, GEP-based receptor-status prediction provided a higher hazard ratio with higher significance in HR– patients (a), implying that GEP-based receptor-status prediction had higher prognostic value than traditional IHC-based HR status characterization. On the other hand, in the hormone-therapy group (b), IHC-based receptor-status characterization was found to be more accurate than GEP-based receptor-status prediction. However, the numbers of samples in the test group (HR- patients) were only 11 and 19 for receptor-status characterization based on IHC and GEP, respectively. Among patients with HR+ non-luminal subtype BRCA (c), IHC-based receptor status had no significant prognostic value, in contrast to GEP-based receptor-status prediction. This finding highlighted that HR+ BRCA patients benefited from hormone therapy, even if they were diagnosed with non-luminal subtype tumors. Among HER2+ patients (d), IHC-based receptor-status characterization exhibited higher prognostic value when considering only the p-value. However, the numbers of patients with IHC-based receptor-status data in the test group (HER2+ patients with TMT) were only 22 and 18 based on IHC and GEP, respectively, and all patients that received TMT survived; hence, the hazard ratio could not be precisely determined (Figure 3 and Table 3). Survival analyses in the METABRIC cohort (excluding patients with a pathological stage of I) showed similar findings, implying that GEP-based receptor-status prediction had higher prognostic significance in terms of patient survival compared to traditional IHC-based receptor-status characterization (Figure 4 and Table 3).

**Figure 3. .**
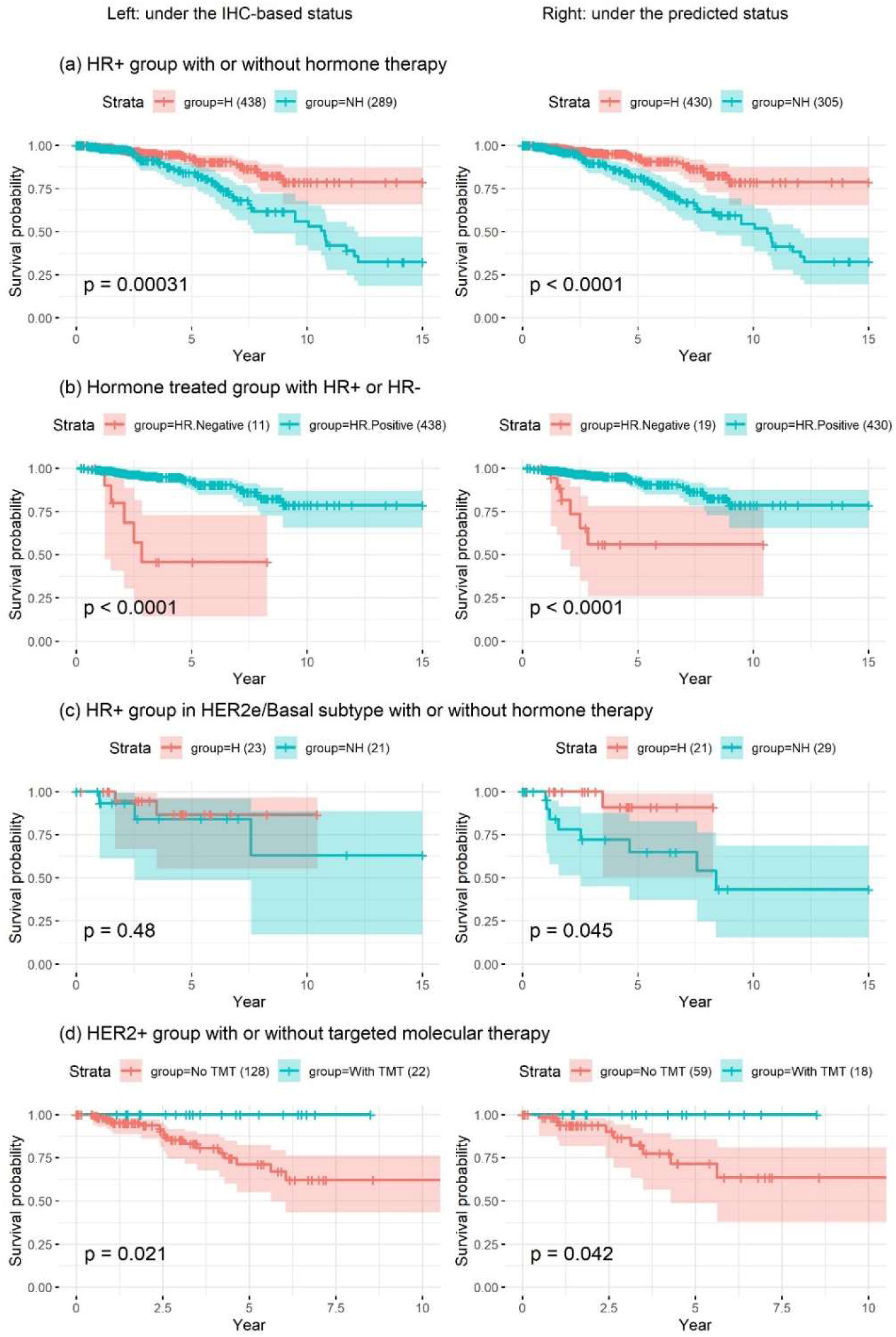
Kaplan–Meier survival analysis of patients from the TCGA dataset using IHC-based (left panel) or GEP-based (right panel) receptor status. Patients were stratified to those who received hormone therapy (H) and those who did not (NH). (a) GEP-based receptor status prediction had higher prognostic significance in terms of patient survival compared to IHC-based HR status. (b) IHC-based receptor-status characterization was found to be more accurate that GEP-based receptor-status prediction. However, the numbers of samples in the test group (HR– patients) for receptor-status characterization based on IHC and GEP were only 11 and 19, respectively. (c) IHC-based receptor status had no significant prognostic value, in contrast to GEP-based receptor-status prediction. (d) The statistical significance of IHC-based receptor-status characterization indicated higher prognostic value. However, the numbers of patients with IHC-based receptor-status data in the test group (HER2+ patients with TMT) were only 22 under IHC and 18 under GEP, and all patients who received TMT survived; hence, the hazard ratio could not be precisely determined.

**Figure 4.**
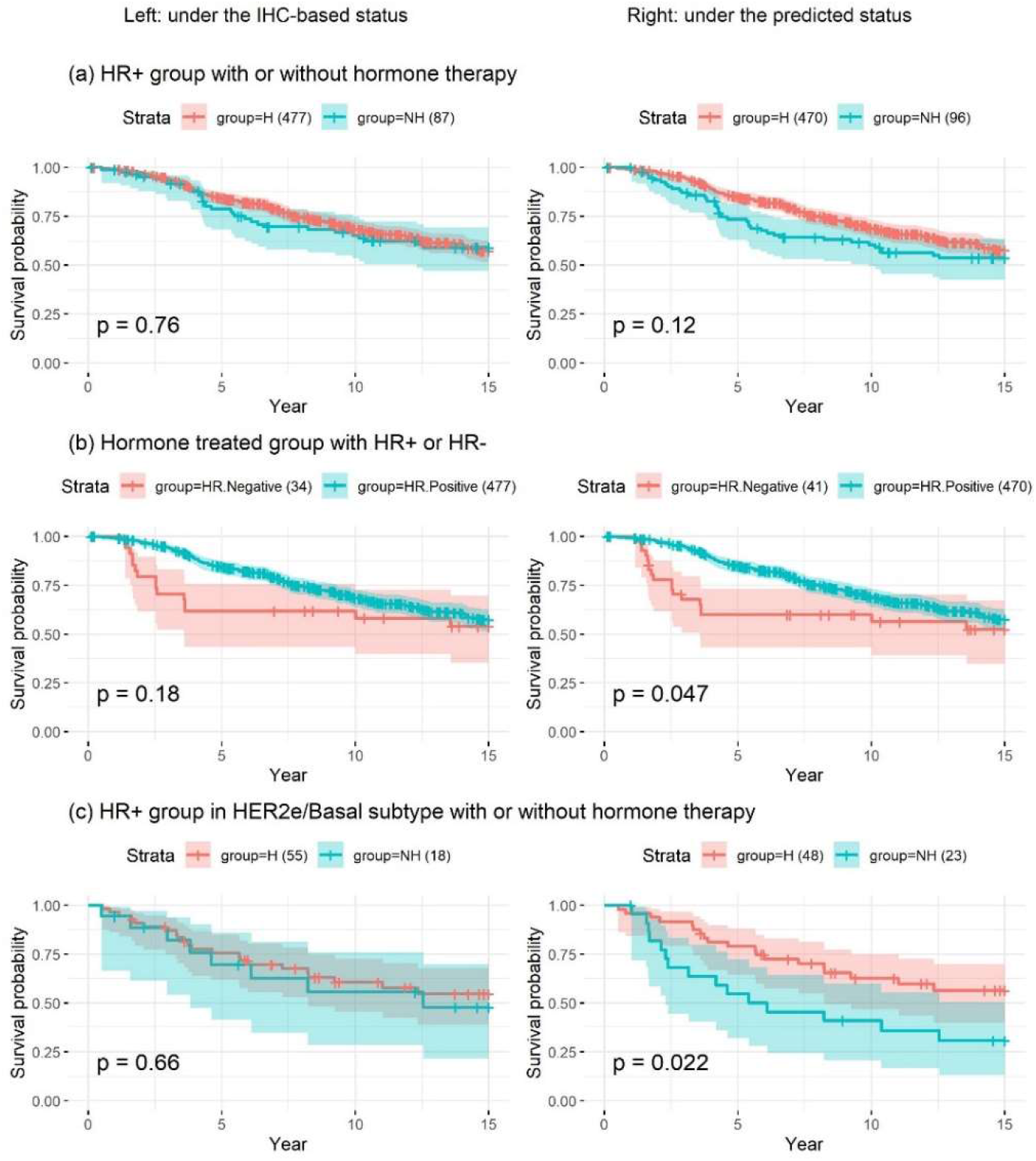
Kaplan–Meier survival analysis in patients of the METABRIC dataset with a pathological stage of II or III (excluding pathological stage I). The analysis was performed using IHC-based receptor status (left panel) or GEP-based receptor status (right panel). GEP-based receptor-status prediction had higher prognostic significance in terms of patient survival compared to traditional IHC-based receptor-status characterization.

**Table 3.** A summary of the hazard ratios and associated statistical significance obtained from survival analyses using IHC-based receptor status (IHC) or the predicted status (pred.). For the survival analysis, data from the TCGA and METABRIC datasets were used.

| Patient group | Conditions compared | # of samples |  | *p*-value |  | Hazard ratio |  |
| --- | --- | --- | --- | --- | --- | --- | --- |
|  |  | IHC | Pred. | IHC | Pred. | IHC | Pred. |
| TCGA |  |  |  |  |  |  |  |
| (a) HR+ | H vs. NH | 727 (438, 289) | 735 (430, 305) | 0.00031 | 2.11·10<sup>-05</sup> | 0.89 | 1.0 |
| (b) Hormone therapy | HR+ vs. HR- | 449 (438, 11) | 449 (430, 19) | 3.15·10<sup>-08</sup> | 3.38·10<sup>-07</sup> | 2.23 | 2.0 |
| (c) HR+ in HER2e/Basal | H vs. NH | 44 (23, 21) | 50 (21, 29) | 0.48 | 0.045 | 0.65 | 1.88 |
| (d) HER2+ | T vs. NT | 150 (22, 128) | 77 (18, 59) | 0.021 | 0.042 | 19.4 | 19.6 |
| METABRIC |  |  |  |  |  |  |  |
| (e) HR+ | H vs. NH | 564 (477, 87) | 566 (470, 96) | 0.76 | 0.12 | 0.06 | 0.28 |
| (f) Hormone therapy | HR+ vs. HR- | 511 (477, 34) | 511 (470, 41) | 0.18 | 0.047 | 0.36 | 0.49 |
| (g) HR+ in HER2e/Basal | H vs. NH | 73 (55, 18) | 71 (48, 23) | 0.66 | 0.022 | 0.18 | 0.77 |

HR: hormone receptor; H: with hormone therapy regardless of chemotherapy; NH: without hormone therapy; T: with targeted molecular therapy regardless of hormone/chemotherapy; NT: without targeted molecular therapy

### 2.5. Patients with non-matching receptor status had significantly worse survival

The type of adjuvant therapy is based mainly on the status of the three receptors. Hence, accurate characterization of receptor status is of high clinical importance. As shown in Figure 5, patients with matching receptor status had longer overall survival (OS) compared to those with non-matching status (hazard ratios 0.6 and 0.79 for the TCGA BRCA and METABRIC cohorts, respectively). Assuming higher accuracy for GEP-based receptor-status prediction, these results highlight the impact of inappropriate treatment due to errors in receptor-status characterization. Although it is unlikely that GEP-based receptor-status prediction is 100% accurate, it can identify patients who can benefit from hormone therapy more reliably than the traditional IHC-based method.

**Figure 5.**
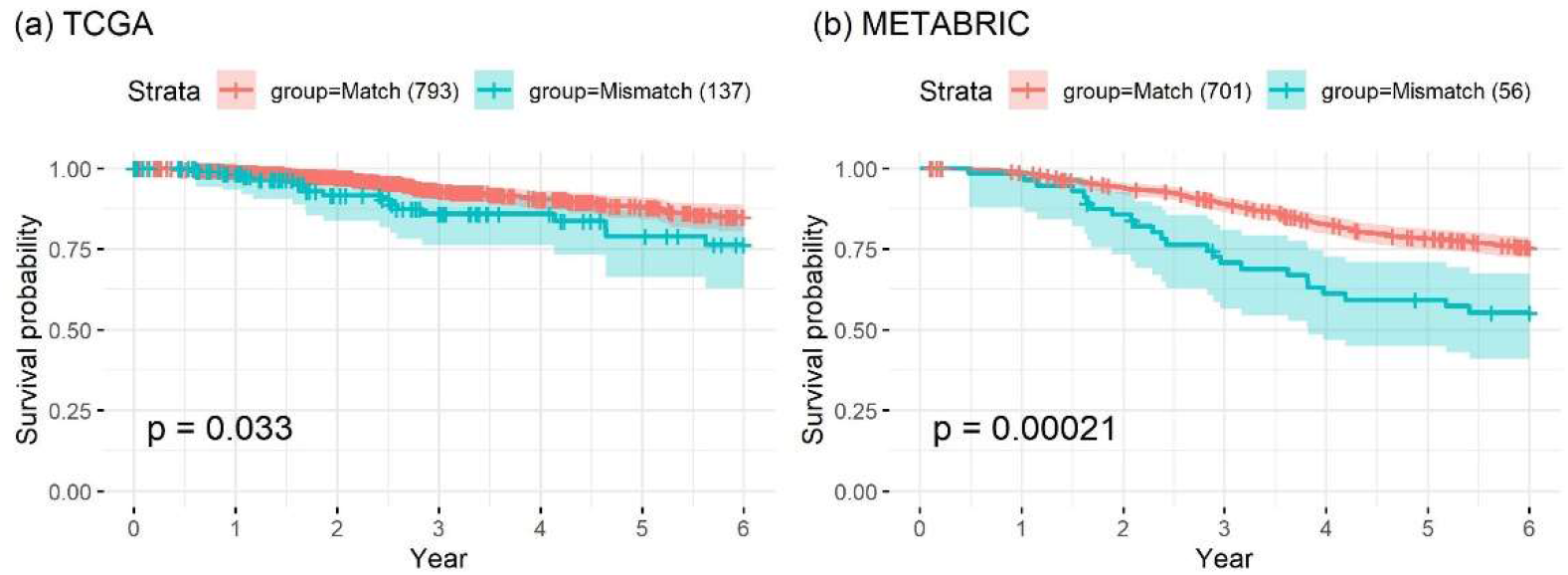
Kaplan–Meier survival analysis of patients in the (a) TCGA BRCA cohort and (b) METABRIC dataset with matching and non-matching receptor status. The hazard ratios of patients with non-matching status were 0.6 for the TCGA BRCA cohort and 0.79 for the METABRIC dataset.

## 3. Discussion

IHC-based assessment of the expression of a specific protein is undoubtedly an important tool for detecting biomarkers in clinical practice. However, this procedure entails severe limitations, including variations in the IHC procedure that can influence the results and their interpretation. As an alternative, biomarker characterization could be performed at the mRNA level; unfortunately, high mRNA levels do not necessarily translate into high levels of the corresponding protein. Additionally, characterization based solely on the expression levels of a single gene or protein inevitably entails the risk of noise. To overcome these limitations, we considered the potential use of GEP-based receptor-status prediction for molecular characterization of BRCA subtypes. Changes in the expression of a gene should be reflected in those of co-expressed genes; therefore, prediction based on the expression of correlated genes may outperform molecular characterization based on a single gene.

In the era of biomarker-assisted targeted therapy, the method used to assess biomarker expression is crucial, as it can improve the prognosis for patients with BRCA and other malignancies. Several challenges remain to be overcome in biomarker-assisted targeted therapies, such as IHC-determined borderline HR-positivity, equivocal HER2 amplification, and discordance between IHC-based subtypes and intrinsic subtypes. Previous studies have shown significant discordance between clinical subtypes and intrinsic subtypes, which affects the prognosis of BRCA patients. Kim et al. reported that discrepancies between the IHC-based subtype and the intrinsic subtype were associated with poor survival, highlighting the limitations of current IHC-based classification methods [30]. Consistent with previous results, we confirmed the poor survival of patients with non-matching subgroup classifications in both the TCGA and METABRIC datasets. These results emphasize the clinical importance of establishing more accurate classification methods. Herein, we evaluated the concordance between the intrinsic subtype and the predicted status of ER, PR, and HER2 using GEP. We found a higher concordance rate between the intrinsic subtype and GEP-based receptor-status prediction compared to receptor status as characterized by IHC. This was consistent in all BRCA subtypes except for the HER2-enriched subtype. These findings imply that GEP-based HR status prediction could be a promising alternative approach to IHC.

Both IHC-based receptor-status characterization and GEP-based status prediction resulted in considerable discordance between HER2-positivity and the HER2-enriched subtype. Although the HER2-enriched subtype is the predominant type of HER2-positive BRCA, three other subtypes exist. A recent study analyzing data from four prospective neoadjuvant trials reported that the percentages of the luminal A, luminal B, HER2-enriched, and basal-like subtypes among HER2-positive BRCA patients were 24%, 20%, 47%, and 9%, respectively [39]. This finding may be partly explained by high intratumoral heterogeneity. Previous genomic analyses have revealed that HER2-positive BRCA is extremely clinically and biologically heterogeneous [40, 41]. The HER2-enriched subtype is also highly heterogeneous, rendering IHC/FISH- and PAM50-based subtyping challenging.

Furthermore, the HER2-enriched subtype can have a distinctive transcriptional landscape independent of HER2 amplification. Analyses in TCGA showed that the HER2-enriched subtype was characterized by the highest number of DNA mutations, including in TP53 and PIK3CA [26]. Recently, Daemen A et al. performed genomic and transcriptomic profiling of HER2-enriched tumors; they concluded that HER2 was not a cancer subtype but rather a pan-cancer phenomenon and that HER2-positive tumors are hormonally driven [42]. Even though further stratification of HER2-enriched BRCA might be beneficial, it might be difficult to achieve further characterization based on GEP. To overcome the limitations of macroscopic GEP, different microscopic prediction approaches could be used, including precise reconstruction of transcriptome data and use of single-cell RNA-seq. These approaches might achieve more in-depth characterization of the molecular subtypes.

To investigate the clinical relevance of GEP-based prediction of ER, PR, and HER2 receptor status, we performed survival analysis of HR+ patients who did or did not receive hormone therapy, as well as of HR+ and HR– patients treated with hormone therapy. GEP-based receptor-status prediction showed a more significant association between treatment outcomes and HR status compared to IHC-based receptor-status characterization. Of note, some benefit was achieved from hormone therapy by patients who were identified as HR+ non-luminal BRCA using GEP-based prediction, in contrast to when IHC-based HR status characterization was performed. These results imply that GEP-based receptor-status prediction can better identify patients who can benefit from hormone therapy, even in patients with non-luminal subtype BRCA. Some studies have shown that adjuvant or palliative hormone therapy is less effective in patients with HR+ BRCA of the non-luminal subtype [43, 44]. However, there is limited evidence regarding which HR+ non-luminal BRCA patients will benefit from hormone therapy. Future studies are needed to determine whether GEP-based receptor-status prediction can address these clinically important questions. In contrast to the HR status, we did not observe improvement in HER2 status prediction; this may be attributed partially to the small number of patients who received targeted molecular therapy for HER2.

## 4. Materials and Methods

The workflow of this study is shown in Figure 6. Our analyses were performed in three steps. First, we identified common predictor genes from two different gene-expression datasets. Second, we predicted ER, PR, and HER2 status based on the shared predictor genes. Finally, we compared survival outcomes according to IHC-based and GEP-based predictions of receptor status.

**Figure 6.**
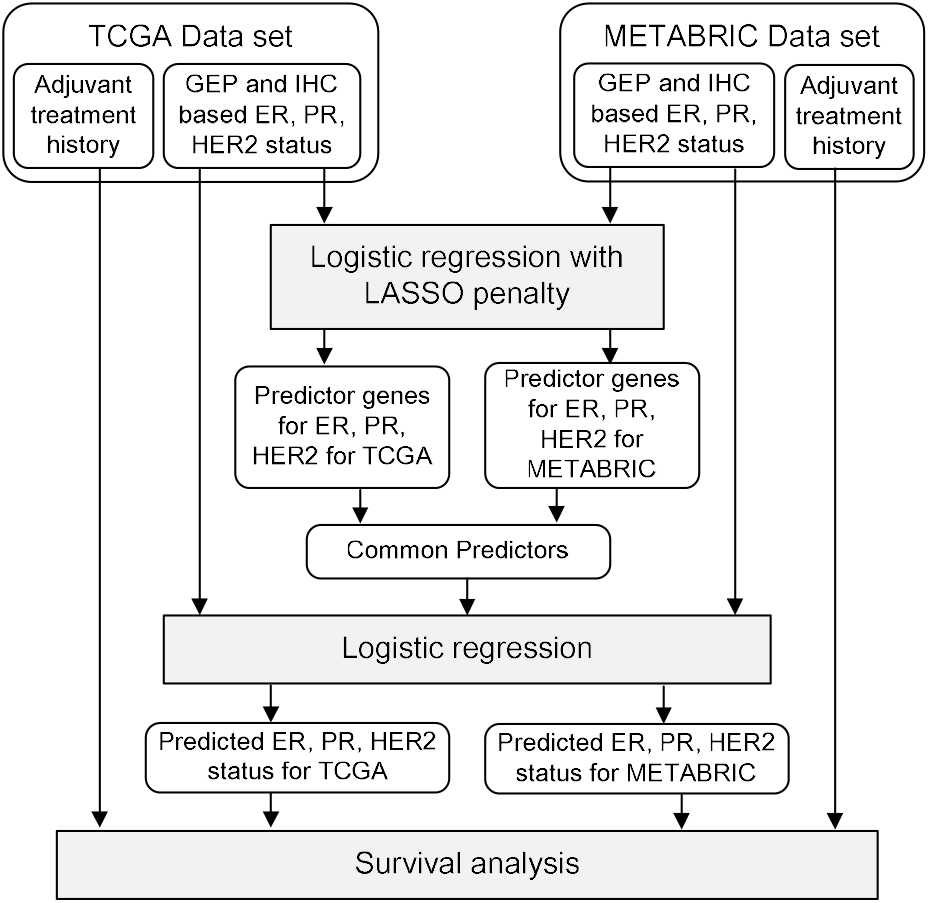
Workflow of gene selection, model training, receptor-status prediction, and survival analysis.

### 4.1. Datasets

For this study, we used BRCA patients’ gene-expression-profile and clinical data acquired from the cancer genome atlas (TCGA) [http://firebrowse.org/] and the Molecular Taxonomy of Breast Cancer International Consortium (METABRIC) databases [https://www.cbioportal.org/] [27]. Both datasets include information on the history of adjuvant treatment, which was a critical element in the survival analyses performed in this study. A summary of the data contained in the two datasets is shown in Table 4.

**Table 4.** A summary of data availability in the TCGA BRCA cohort and METABRIC dataset.

| Item | TCGA BRCA cohort | METABRIC | Comment |
| --- | --- | --- | --- |
| Gene expression profile | Yes | Yes |  |
| PAM50-based subtype | Yes (partially) | Yes |  |
| ER status | Yes (IHC) | Yes (IHC, non-IHC) | Used IHC-based status |
| PR status | Yes (IHC) | Yes (non-IHC) | Used for receptor status |
| HER2 status | Yes (IHC) | Yes (IHC, non-IHC) | Used IHC-based status |
| RPPA measurements | Yes | No |  |
| Types of drug treatment | Chemo, hormone and targeted molecular therapy | Chemo and hormone therapy | Used for survival analysis |
| Age at initial diagnosis | Yes | Yes | Used for sample selection |
| Pathological stage | Yes | Yes | Used for sample selection |

The TCGA BRCA dataset contained data from tumor samples (n = 1,092 patients) and adjacent normal tissues (n = 112 patients). The METABRIC dataset contained data from 2,506 tumor samples, including GEP data from 1,904 patients. The TCGA and METABRIC datasets also contained clinical data, including ER, PR, and HER2 status, as well as histories of surgery, radiation-therapy, and drug treatments; however, clinical data were not available for all of the patients. Information regarding the tumor subtype was available for some samples in the TCGA BRCA dataset; PAM50 mRNA profile information was available for 523 of 1,092 patients [26]. To ensure consistency between the two datasets, information on ER and HER2 status as determined by IHC was used for patients in the METABRIC dataset. Non-IHC-based PR status was used for the METABRIC cohort because the PR status was not assessed by IHC in these patients.

### 4.2. Prediction model and gene selection

Based on GEP and the status of the three receptors, logistic regression with LASSO penalty was performed in a supervised mode to identify predictor genes for each of the two datasets. This analysis was performed using the R package glmnet [45–47]. In the TCGA BRCA dataset, the expression levels of 17,202 genes were log2-transformed and normalized. In the METABRIC dataset, already normalized mRNA expression data were used. To identify the common predictor genes and minimize overfitting-related errors, LASSO penalty weights were selected for a set of predefined genes (e.g., 10, 20, 40, and 60), and for each number, the penalty weight that led to the closest number of selected genes was chosen. This approach was conducted separately for the TCGA and METABRIC datasets. Common predictor genes between TCGA and METABRIC were then identified to avoid dataset-related dependencies. After inspecting the overall number of shared genes, 40 genes were selected; these contained 7, 6, and 4 common predictor genes for ER, PR, and HER2, respectively, as summarized in Table 1. Subsequently, logistic regression was performed again to train the models for ER, PR, and HER2 status prediction for both TCGA and METABRIC. The mismatch rate was obtained by fivefold cross-validation.

Pairwise correlations of gene-expression levels between the selected genes are shown in Supplementary Figures 4, 5, and 6. Of note, PR predictor genes included *ESR1* and *AGR3*, which were also ER predictor genes. Furthermore, among the four HER2 predictor genes, *CPB1*, *GSTT1*, and *PROM1* showed only small correlations with ERBB2, implying that HER2 status prediction was determined predominantly by ERBB2.

### 4.3. Survival analysis for accuracy evaluation and sample selection

The survival analyses were performed for various group/condition pairs; significance (p-value) was used as an accuracy criterion. Cox’s proportional hazard model was used to determine overall survival [48]; the analysis was repeated using the IHC-based status and the predicted status. For the survival analysis based on IHC-based receptor status, we used those samples for which IHC-based receptor status was available. For the survival analysis based on predicted-receptor status, we used the same set of samples without considering discrepancies between the predicted status and the IHC-based status. As shown in Table 1, in 5–12% of cases, the predicted status differed from the IHC-based status.

Additionally, for the survival analyses, patients were selected according to the following criteria: (1) pathological cancer stage I, II, or III and (2) age <80 years at initial diagnosis. Subsequently, patients were stratified according to adjuvant drug treatments. The characteristics of the patients included in the survival analyses are summarized in Table 5.

**Table 5.** A summary of the samples available in the TCGA and METABRIC datasets.

| Variable | Conditions | The number of available samples |  |
| --- | --- | --- | --- |
|  |  | In TCGA | In METABRIC |
| Age | ≤80 years | 1,039 | 1,783 |
| Pathologic stage: | I | 170 | 464 |
|  | II | 598 | 736 |
|  | III | 232 | 105 |
| Therapy applied: | Chemotherapy | 578 | 393 |
|  | Hormone therapy | 495 | 1,084 |
|  | Both chemo- and hormone therapy | 324 | 181 |
|  | Targeted molecular therapy | 30 | NA |
| ER status: | Positive | 760 | 1,339 |
|  | Negative | 230 | 418 |
|  | NA | 2 | 0 |
| PR status: | Positive | 663 | 946 |
|  | Negative | 324 | 837 |
|  | NA | 4 | 0 |
| HER2 status: | Positive | 159 | 114 |
|  | Negative | 524 | 647 |
|  | NA | 182 | 27 |

\* For ER, PR, and HER2 status; 'indeterminate' and 'equivocal' were reported as NA.

## 5. Conclusions

Therapeutic decision making in BRCA is heavily based on the clinical subtype defined by HR and HER2 expression status. NGS-based approaches could allow more accurate characterization of the various molecular and clinical features of BRCA. GEP-based receptor-status prediction could provide a better understanding of BRCA pathology and guide physicians in decision making. To improve the performance of GEP-based prediction models, data from larger cohorts are required for standardization of the procedure. In addition, a more comprehensive analysis of receptor status should be performed to identify the characteristics that affect the positivity or negativity of the status of the three receptors, as well as the mechanisms responsible for the discordance between intrinsic subtype and clinical subtype.

## Supplementary Materials

The following materials contain some of TCGA and METABRIC clinical data and the new predictions on the 3-receptor status, which were used for the survival analyses in this work.

1. TCGA_BRAC_clinical_data_n_pred_status.csv:
2. METABRIC_clinical_data_n_pred_status.csv

## Author Contributions

Conceptualization, S.Y., H.S.W., K.K., W.J.P. and Y.H.K.; methodology, S.Y., K.K. and W.J.P., software, validation and formal analysis, S.Y., and K.Q.; investigation and data curation, H.S.W. and Y.H.K.; funding acquisition, S.Y.; writing—original draft preparation and visualization, S.Y., H.S.W and K.K., writing—review and editing, S.Y., H.S.W., K.K., W.J.P. and Y.H.K.; All authors have read and agreed to the published version of the manuscript.

## Acknowledgments

The results here are in part based upon data generated by the TCGA Research Network: https://www.cancer.gov/tcga.

## Supplemental Figures

**Supplemental Figure 1.**
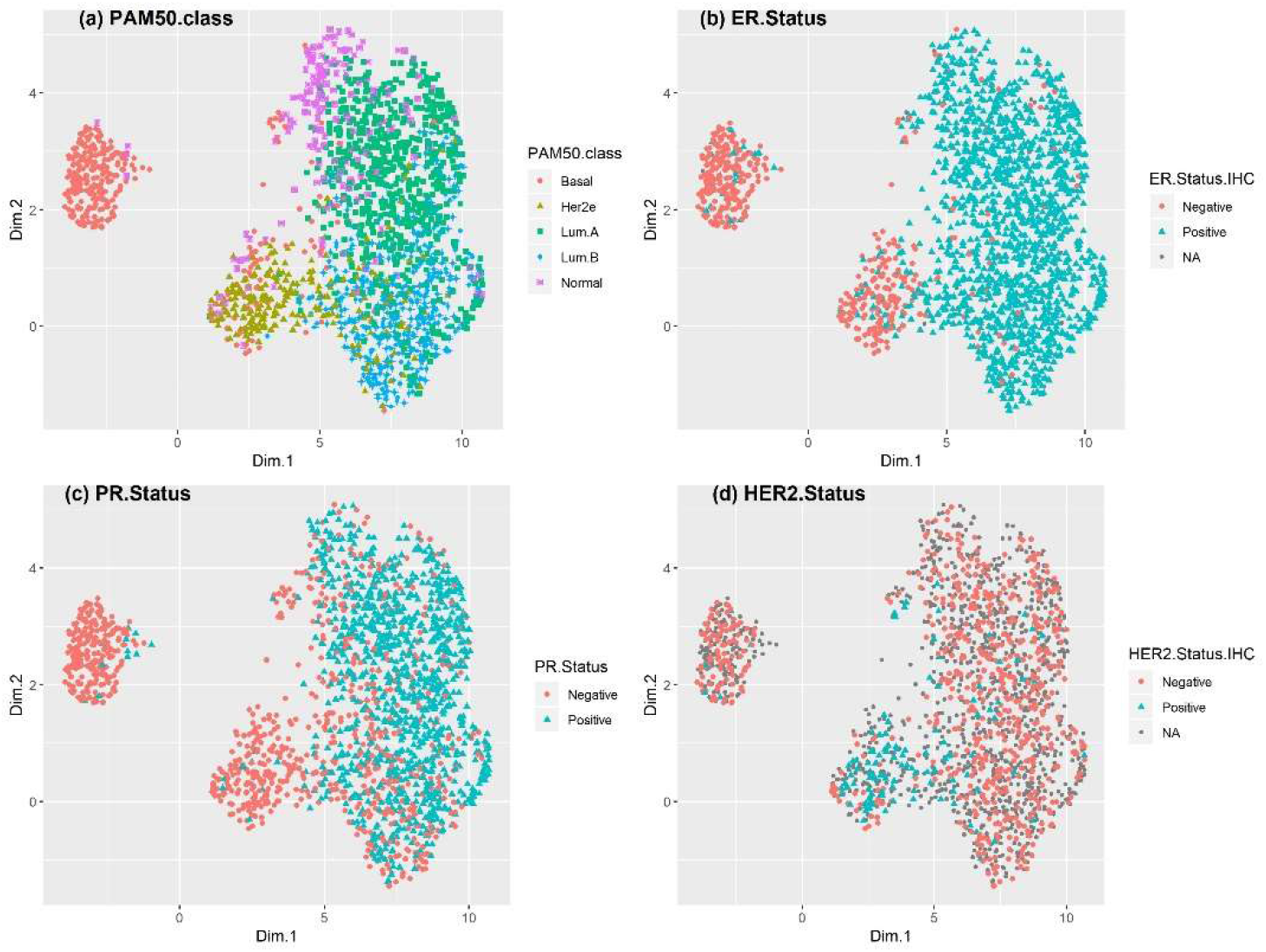
UMAP plot showing receptor status of patients in the METABRIC dataset. The tumor subtype and ER, PR, and HER2 status were based on the available clinical data. Gray points are samples with no available clinical information. The UMAP plot of the METABRIC dataset revealed a similar macroscopic landscape to that for TCGA.

**Supplemental Figure 2.**
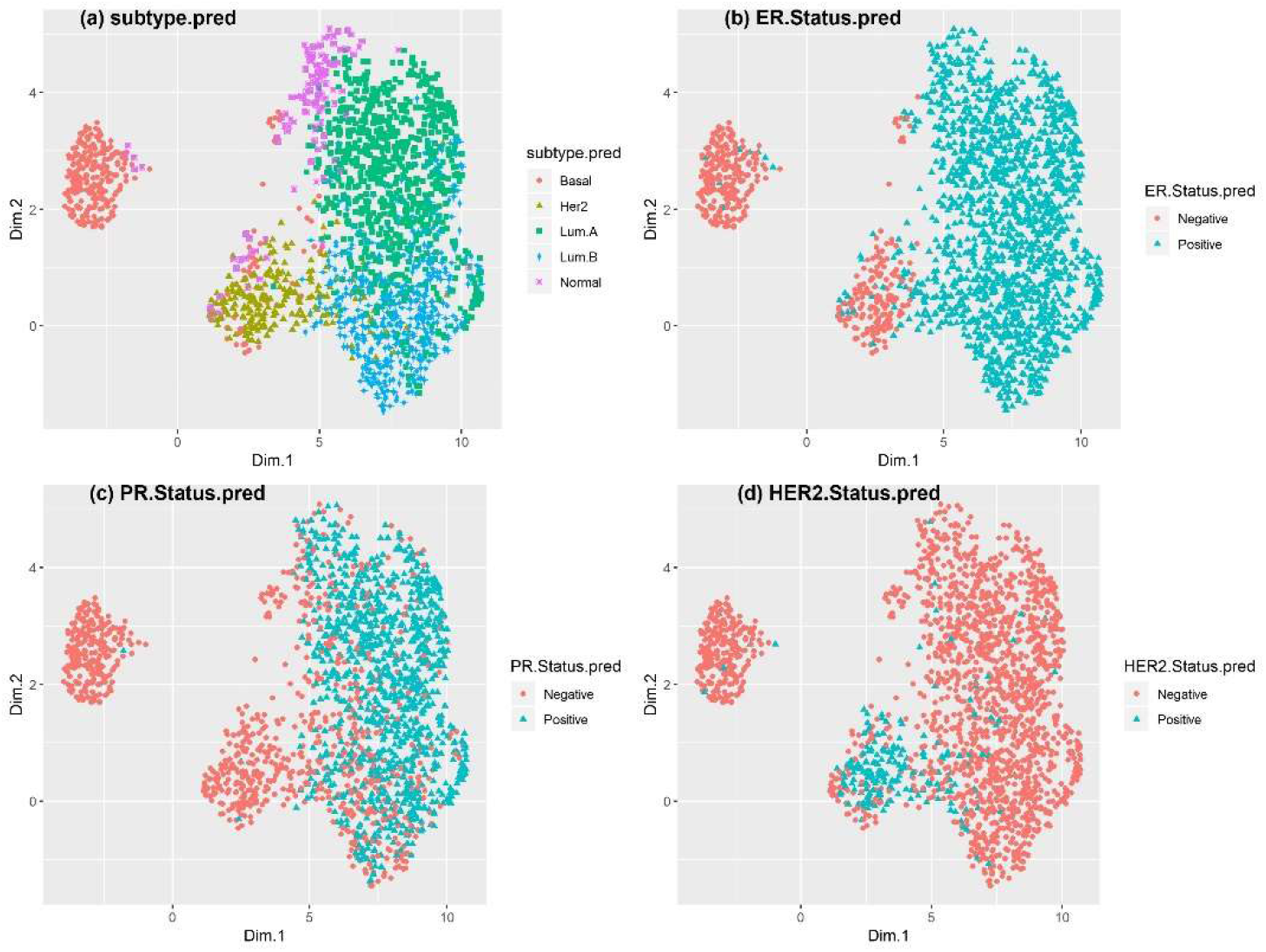
UMAP plot showing receptor status of patients in the METABRIC dataset. GEP-based prediction was used to determine the subtype, as well as the status of ER, PR, and HER2. Similar to TCGA, the predicted ER and HER2 status (but not PR) was mostly in accordance with the corresponding pattern of receptor status for the basal-like, luminal A, and luminal B subtypes.

**Supplemental Figure 3.**
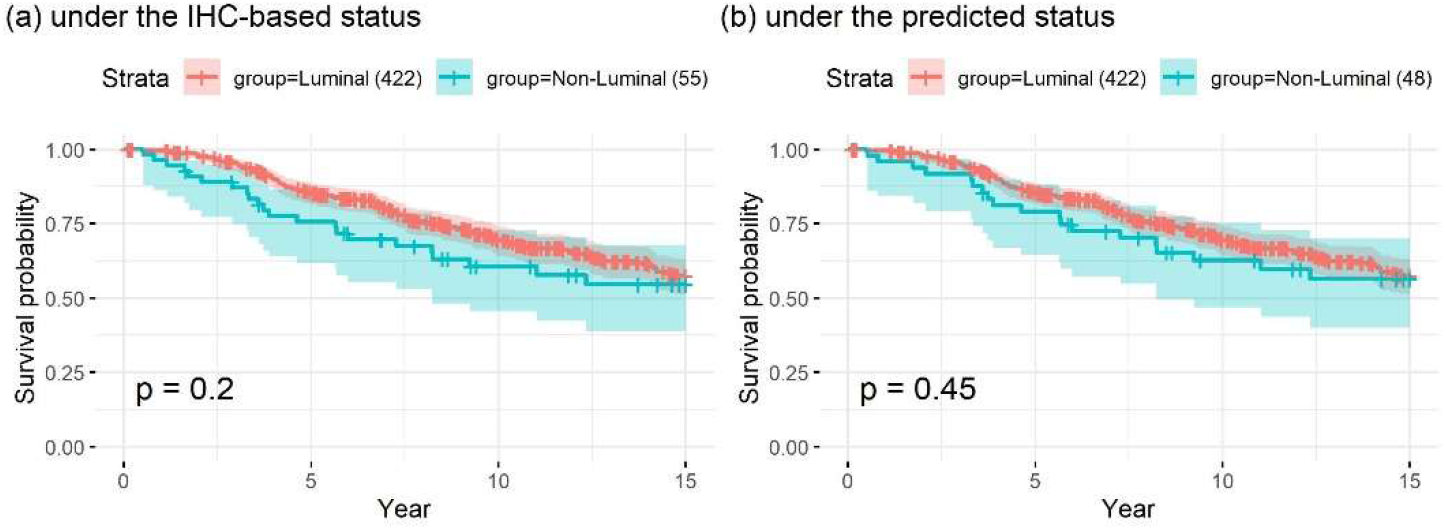
Scatter plots and Pearson’s correlation coefficients of seven predictor genes for ER status prediction. Blue: ER+; red: ER−; empty circle: NA. ER status characterization was based on IHC.

**Supplemental Figure 4.**
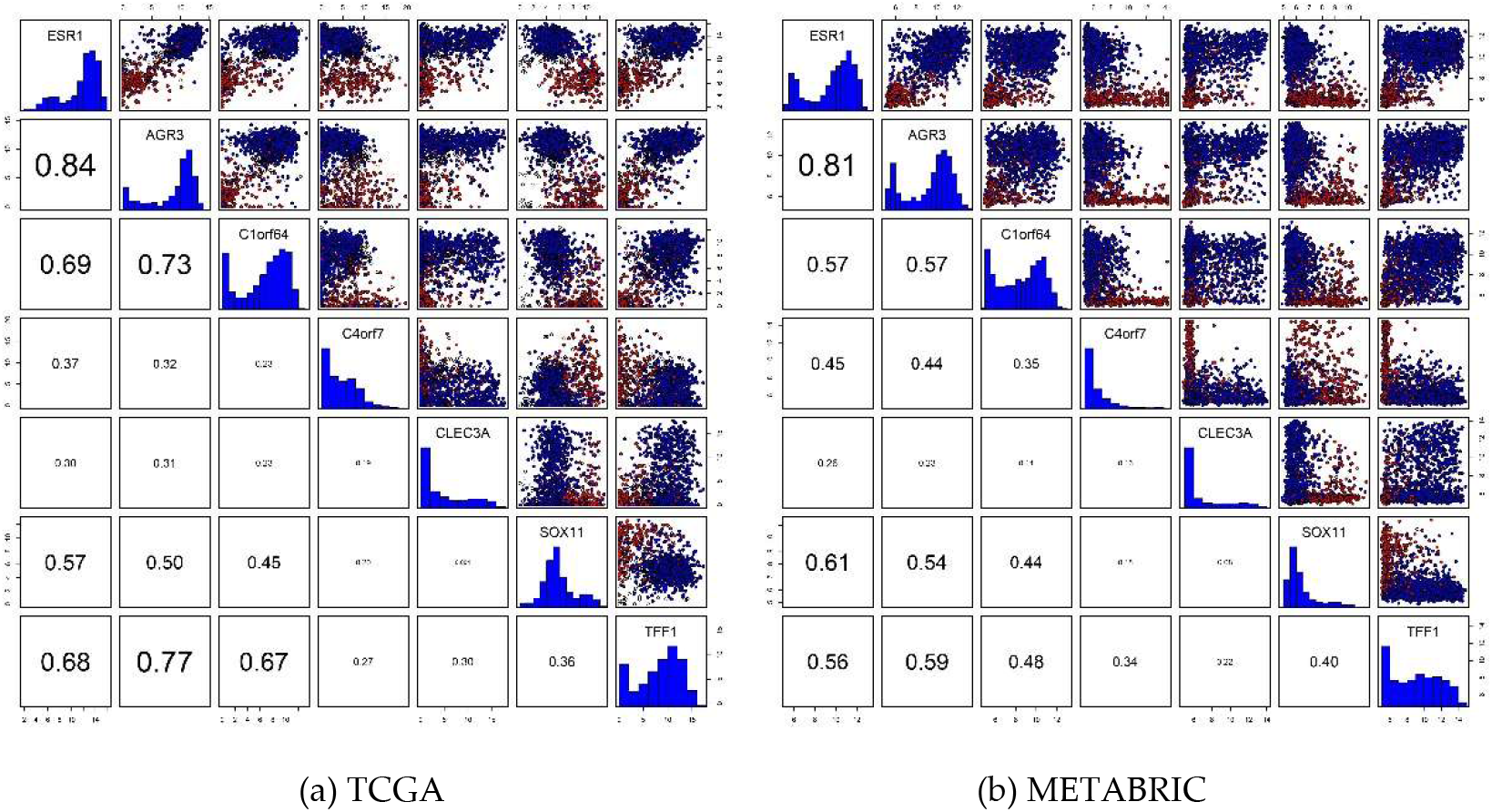
Scatter plots and Pearson’s correlation coefficients of seven predictor genes for ER status prediction. Blue: ER+; red: ER−; empty circle: NA. ER status characterization was based on IHC.

**Supplemental Figure 5.**
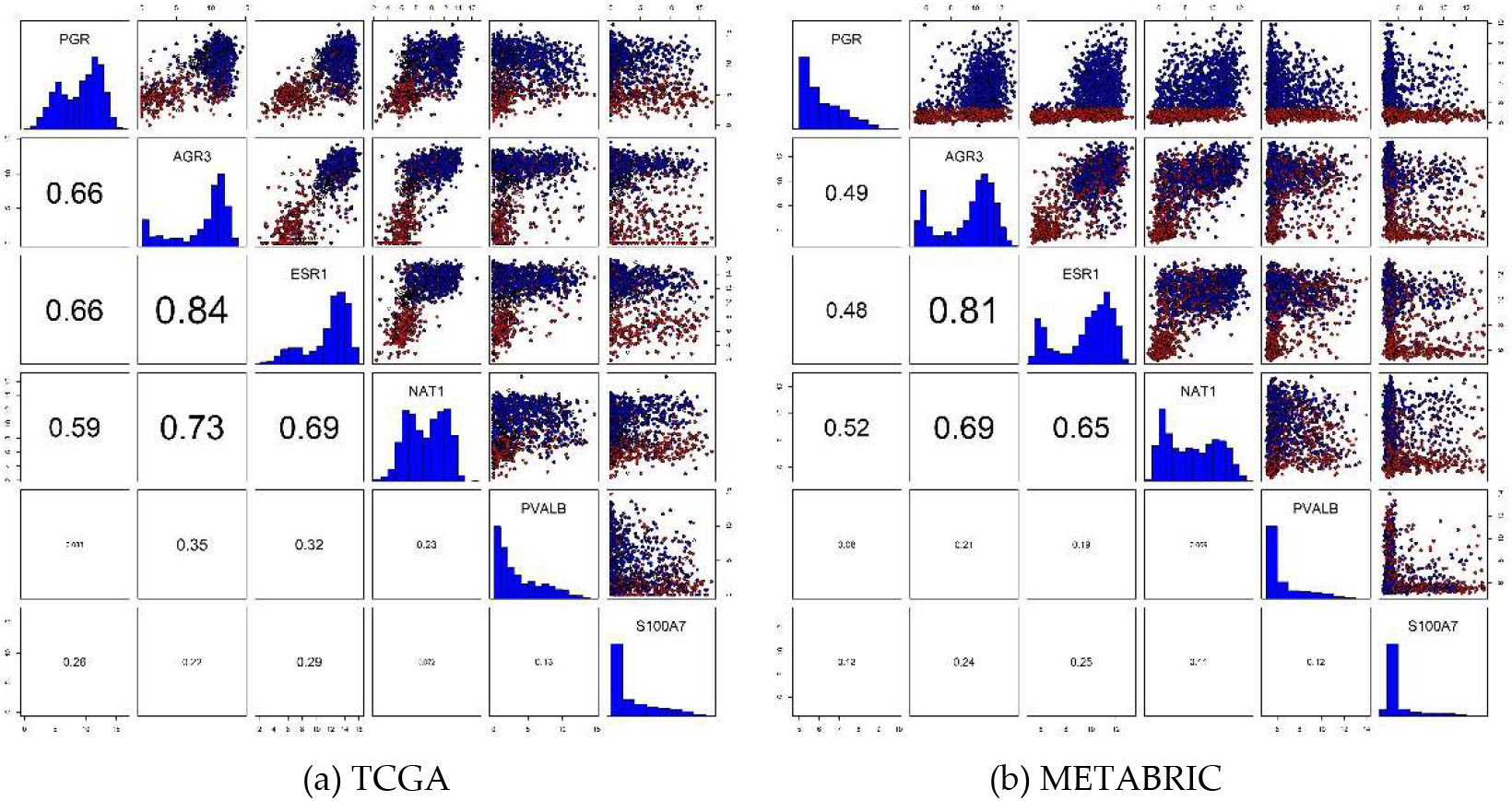
Scatter plots and Pearson’s correlation coefficients of six predictor genes for PR status prediction. Blue: PR+; red: PR−; empty circle: NA. The PR status of TCGA samples was based on IHC, whereas that of METABRIC samples was not based on IHC. The PR-status predictor genes included *ESR1* and *AGR3*, which were also predictor genes for ER status.

**Supplemental Figure 6.**
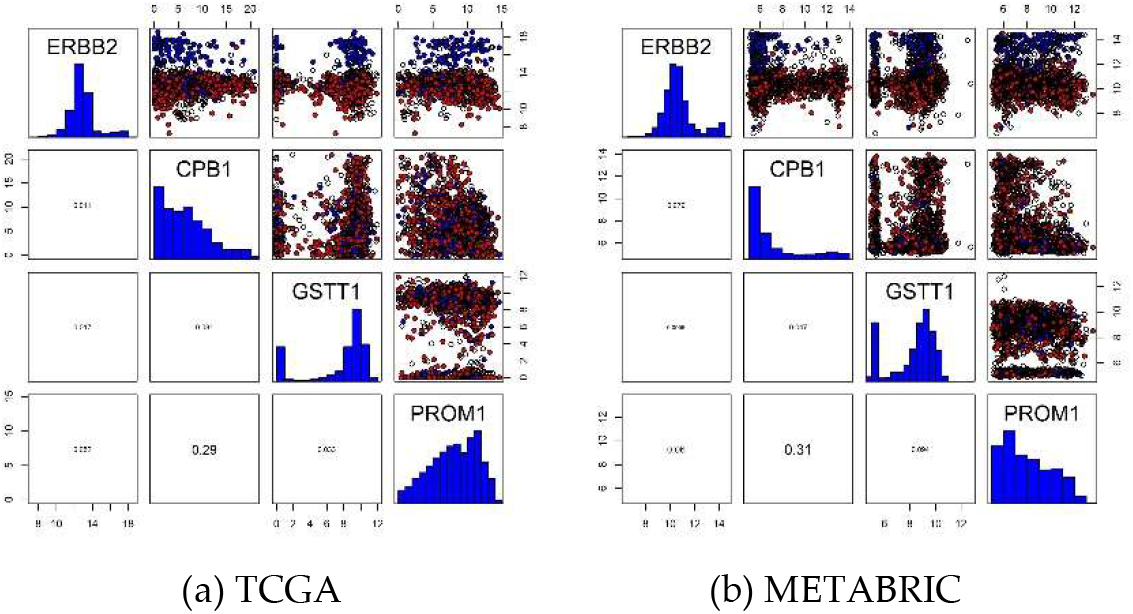
Scatter plots and Pearson’s correlation coefficients of four predictor genes for HER2 status prediction. Blue: HER2+; red: HER2−; empty circle: NA. *CPB1*, *GSTT1*, and *PROM1* showed weak correlations with *ERBB2*, implying that HER2 status prediction was determined predominantly by *ERBB2*.

